# Why Is Spontaneous Blink Timing Informative? An Adaptive Scheduling Perspective

**DOI:** 10.64898/2026.09.21.753188

**Authors:** Stefan Arnau, Anna Plachti, Emad Alyan, Edmund Wascher, Daniel Schneider

**Affiliations:** Leibniz Research Centre for Working Environment and Human Factors Dortmund (IfADo), Ardeystraße 67, 44139 Dortmund, Germany

**Keywords:** Spontaneous eye blinks, Blink timing, Mental chronometry, Visual sampling, Neuroergonomics

## Abstract

Spontaneous eye blinks have long been linked to cognitive processing, yet how task demands shape blink timing and its relationship to behavioral performance remains unclear. We examined spontaneous blink behavior in 576 adults performing two variants of the Continuous Performance Task (CPT). Blink occurrence and timing were most strongly modulated by the experimental condition in the more demanding CPT-AX task, whereas their association with response time was stronger in the CPT-X task, where more consistent blink timing predicted faster responses. This dissociation suggests that task structure changes not only blink behavior but also the behavioral relevance of blink timing. These findings are consistent with an adaptive scheduling account of spontaneous blinking and provide a conceptual framework for understanding when and why blink timing contains chronometric information about ongoing cognition.

## Introduction

Understanding how cognitive processing is organized over time has long been a central objective of mental chronometry and cognitive neuroscience (Meyer et al., 1988; Posner, 2005). Many established methods provide valuable insights into this temporal organization, including electroencephalography (EEG), functional neuroimaging, and eye tracking. However, these approaches often require specialized equipment, substantial analytical effort, or laboratory-based recording environments (Wascher et al., 2023).

Consequently, there is continued interest in identifying behavioral measures that are simple to acquire, temporally precise, and informative about the temporal organization of cognitive processing.

Spontaneous eye blinks have recently emerged as one such candidate. Although blinking is primarily required to maintain the corneal tear film, spontaneous blinks occur considerably more frequently than is necessary for lubrication (Kaminer et al., 2011; Zametkin et al., 1979) and are systematically influenced by cognitive state (Chermahini & Hommel, 2010; Colzato et al., 2008). Across a broad range of experimental paradigms, blink behavior is systematically modulated by attentional allocation (Bonfiglio et al., 2011; Maffei & Angrilli, 2018; Oh et al., 2012; Wascher et al., 2015), task demands (Benedetto et al., 2011; Fairclough et al., 2005; Recarte et al., 2008), and task engagement (Fairclough & Venables, 2006). These findings suggest that spontaneous blinking is closely coupled to cognitive activity rather than reflecting a purely physiological maintenance process.

Recent work suggests that the temporal organization of blinking conveys substantially richer information than overall blink frequency. Blink rate is affected by numerous factors, including fatigue, arousal, task characteristics, and pronounced interindividual variability, making it difficult to interpret as a specific marker of cognition (Nakano & Miyazaki, 2019; Stern et al., 1984; Wascher et al., 2015). In contrast, the timing of individual blinks exhibits systematic relationships with meaningful events during task performance. Blinks preferentially occur around sentence endings during reading (Cornelis et al., 2025; Orchard & Stern, 1991), after decisions have been completed (Fukuda, 2001), at narrative event boundaries during movie viewing (Nakano et al., 2009), during speech perception (Kobald et al., 2019), and following periods of intensive stimulus evaluation in laboratory tasks (Wascher et al., 2015). Likewise, reproducible blink timing patterns have been observed during naturalistic activities such as walking or Formula car driving, where blink probability varies systematically with momentary task demands (Nishizono et al., 2023; Wascher et al., 2022)

Collectively, these findings suggest that the temporal distribution of spontaneous eye blinks is systematically related to the temporal organization of cognitive processing. Although a growing body of evidence supports this relationship, the mechanisms underlying it remain poorly understood (Nakano & Miyazaki, 2019; Nishizono et al., 2023; Wascher et al., 2015). Current evidence therefore supports blink timing as an informative correlate of cognitive processing. However, whether spontaneous blinks directly reflect the completion of processing episodes and transitions between cognitive states, or instead arise for other functional reasons, remains an open question.

An alternative interpretation emerges when blink timing is viewed from a neuroergonomic perspective. Every spontaneous blink transiently interrupts visual input and therefore entails a temporary loss of visual information (Casse et al., 2007; Doughty, 2001; Kwon et al., 2013; Pitigoi et al., 2024). At the same time, regular blinking is physiologically indispensable and cannot simply be suppressed indefinitely (Al-Abdulmunem, 1999). Unlike many physiological processes, spontaneous blinks exhibit considerable temporal flexibility. This creates a scheduling problem: because blinking is both necessary and temporarily costly, blinks should ideally occur when their expected informational cost is minimal (Hoppe et al., 2018; Murali & Händel, 2021; Shultz et al., 2011). Moments following the completion of cognitive processing are likely to represent such opportunities because they coincide with reduced demands for immediate visual sampling. Consequently, the temporal alignment between blinking and cognitive event boundaries does not necessarily imply that blinks directly reflect transitions between cognitive processes. Instead, blink timing may emerge from adaptive scheduling that minimizes transient information loss while simultaneously satisfying physiological requirements.

These perspectives are not mutually exclusive but operate at different explanatory levels. The cognitive account emphasizes the temporal organization of internal information processing, whereas the neuroergonomic perspective emphasizes the adaptive scheduling of visual information sampling. Under these perspectives, blink timing may either directly reflect transitions between successive processing episodes or represent an epiphenomenon arising because both are shaped by the same temporal structure of task demands. Despite these different interpretations, both perspectives converge on the same empirical prediction: blink timing should provide information about ongoing cognitive processing. If blink timing captures aspects of the temporal organization of cognitive processing, and if this organization contributes to successful task performance, blink timing should likewise be systematically related to behavioral performance.

Evaluating the predictive properties of blink timing requires a task context in which the temporal organization of cognitive processing and the temporal structure of task demands are both well defined. The Continuous Performance Task (CPT) (Sharifian et al., 2021; Walter et al., 1964) provides such a context. Each trial consists of a cue followed by a target after a fixed interval. During this cue-target interval, cue information is evaluated, task-relevant representations are established and maintained, expectations about the upcoming target are formed, and preparatory response processes unfold (Braver, 2012; Corbetta & Shulman, 2002). Because the cue must be sampled while behaviorally relevant visual information is expected at the upcoming target, the timing of a transient visual interruption may also have consequences for subsequent performance. These characteristics make the cue-target interval particularly well suited for examining how blink timing relates to the temporal organization of cognitive processing and subsequent behavior.

The present study used data from the Dortmund Vital Study, a large adult cohort study (Gajewski et al., 2023), and employed two variants of the CPT, in which spontaneous blinking could be examined during a fixed cue-target interval preceding target onset.

Across the four task-by-cue conditions, predictive cues systematically varied the behavioral significance of cue information and, consequently, the cognitive demands associated with cue evaluation, target expectation, and response preparation. These manipulations systematically altered both the temporal organization of cognitive processing, and the expected informational costs associated with transiently interrupting visual information sampling. Although the temporal structure of the cue-target interval remained identical across conditions, the temporal profile of these demands differed systematically. The combination of systematic experimental manipulation and a large cohort provides a controlled and well-powered framework for characterizing blink timing across individuals, testing its adaptation to changing task demands, and evaluating its behavioral relevance.

We therefore examined whether spontaneous blink timing during the cue–target interval can serve as a useful chronometric measure of the temporal organization of cognitive processing. The analysis focused on the timing of the first spontaneous blink because previous work has suggested that it provides a more sensitive measure of task-related blink timing than aggregate measures such as overall blink rate (Ichikawa & Ohira, 2004; Wascher et al., 2015). Figure 1 provides an overview of spontaneous blink behavior across the trial cycle, whereas Figure 2 summarizes the blink measures derived from the cue–target interval and their descriptive characteristics. First, blink occurrence, first-blink timing, and blink-timing variability were examined for systematic adaptation to experimentally manipulated task demands. Second, stable individual differences in these blink characteristics were tested for association with subsequent behavioral performance. Finally, trial-to-trial deviations from an individual’s characteristic blink timing were likewise evaluated for their association with behavioral performance.

**Figure 1.**
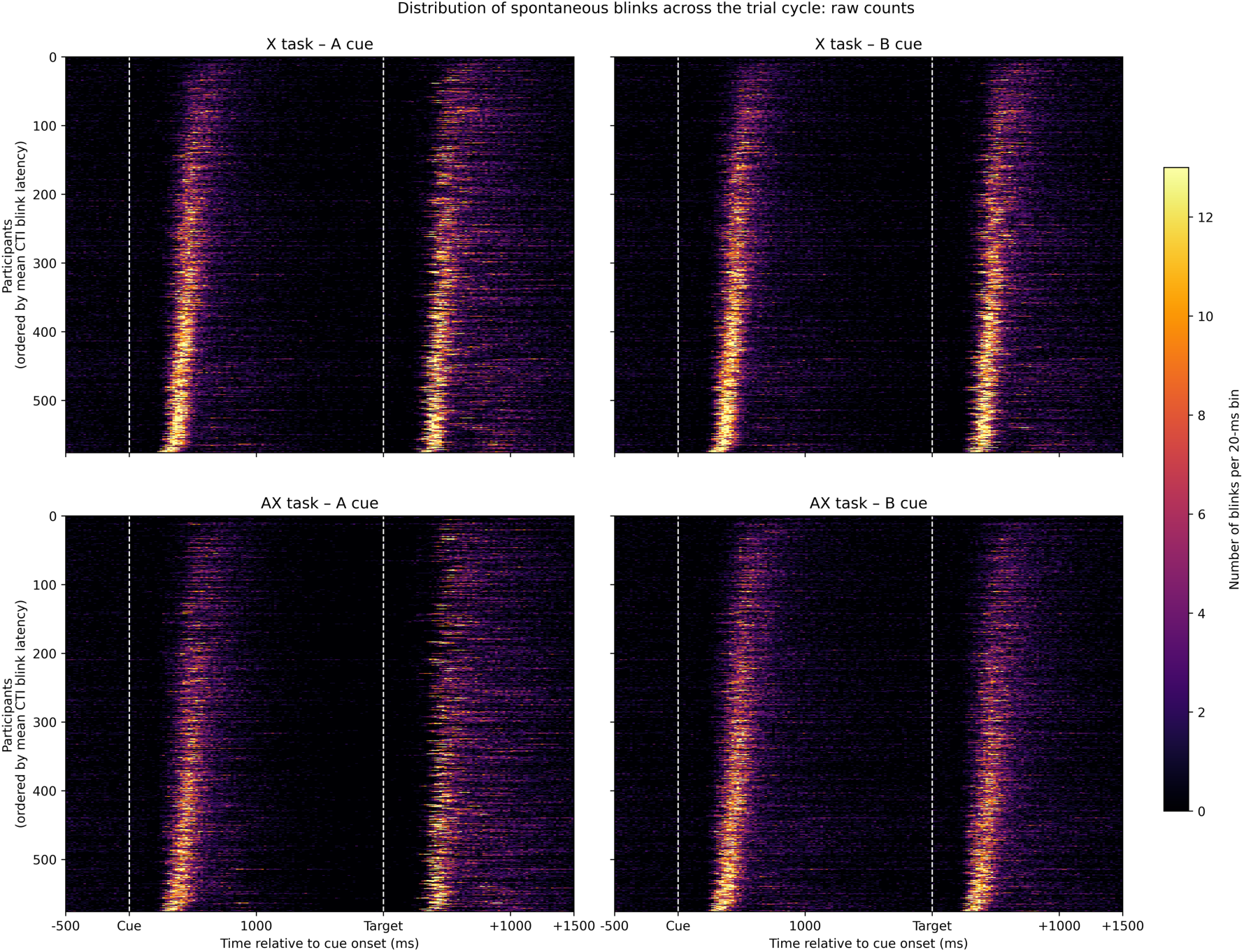
Temporal distribution of spontaneous eye blinks across the trial cycle. Heatmaps show the distribution of spontaneous eye blinks across the complete trial cycle for each combination of task (X, AX) and cue (A, B). Each row represents one participant, ordered by mean cue–target interval (CTI) blink latency. Color indicates the number of blinks within successive 20-ms time bins across all trials. Vertical dashed lines indicate cue onset (0 ms) and target onset (2000 ms).

**Figure 2.**
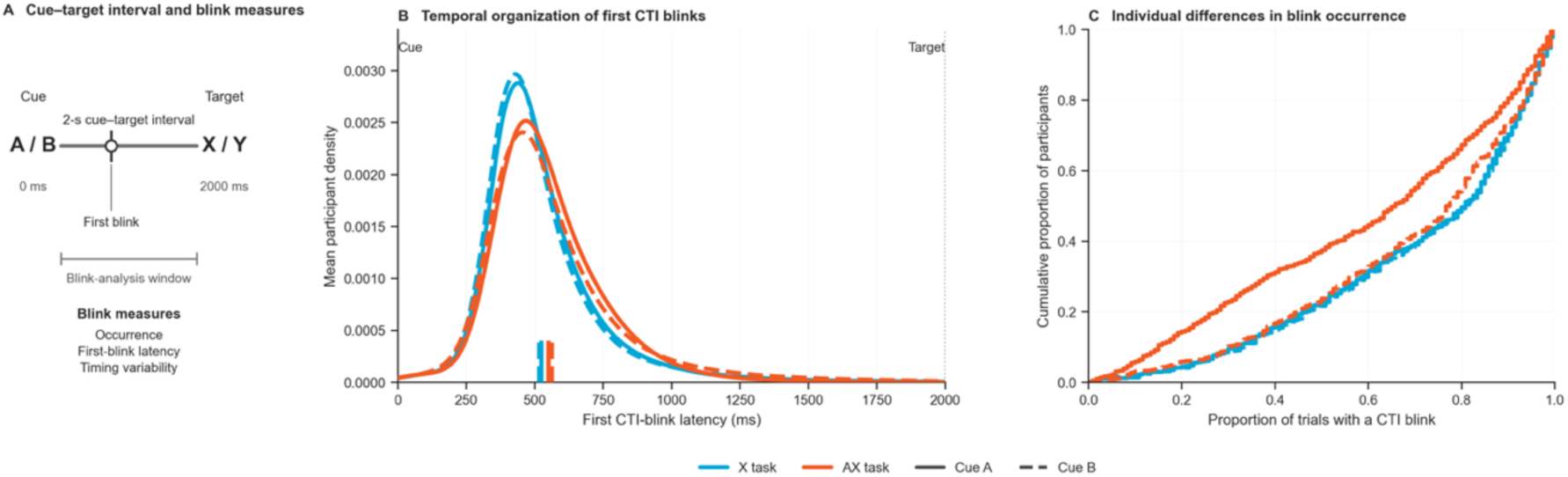
Cue–target interval and descriptive characteristics of spontaneous blink behavior. The figure illustrates the cue–target interval (CTI), the blink measures derived from this interval, and the distributions of blink timing and blink occurrence across task conditions. **(A)** Schematic of the CTI and the blink measures derived from this interval: blink occurrence, first-blink latency, and blink-timing variability. **(B)** Mean participant-level kernel density estimates of first CTI-blink latency, shown separately for the X (blue) and AX (orange) tasks and Cue A (solid) and Cue B (dashed). Short vertical markers indicate participant-level mean first-blink latencies. **(C)** Empirical cumulative distribution functions of participant-level blink occurrence for each Task × Cue condition. At each value on the x-axis, the curve indicates the proportion of participants with blink occurrence at or below that value.

Together, these analyses evaluate whether spontaneous blink timing not only adapts systematically to changing task demands but also provides a behaviorally relevant chronometric measure of the temporal organization of cognitive processing.

## Results

### Task context systematically modulates spontaneous blink behavior

Spontaneous blink behavior was first examined for systematic adaptation to experimentally manipulated task demands. Figure 3 summarizes participant-level distributions, condition means, and mixed-effects model estimates for blink occurrence, first-blink latency, and blink-timing variability across Task × Cue conditions.

**Figure 3.**
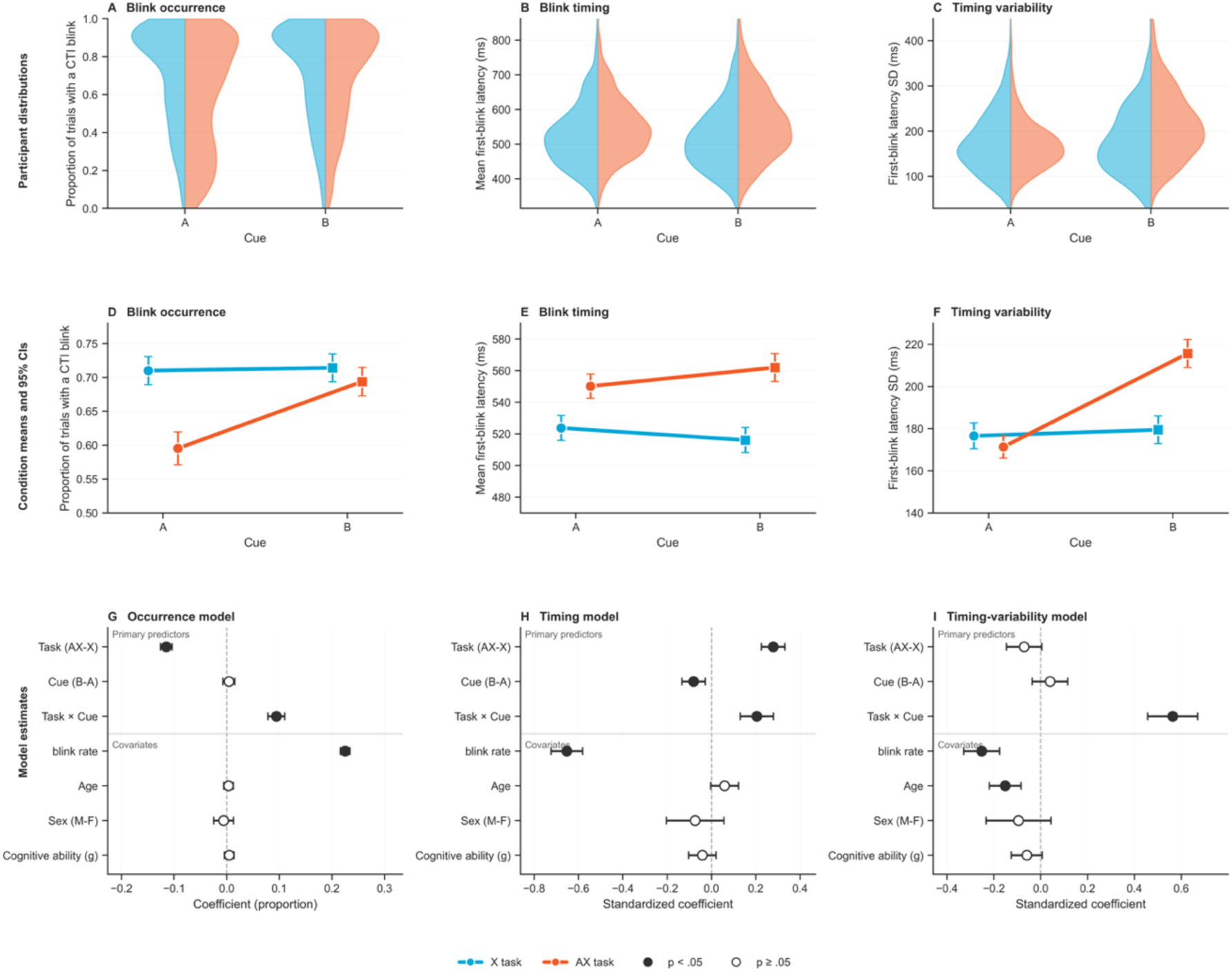
Blink behavior as a function of task context. The figure summarizes participant-level distributions, condition means, and model estimates for blink occurrence, first-blink latency, and blink-timing variability across task and cue conditions. **(A–C)** Participant-level distributions of blink occurrence (A), mean first-blink latency (B), and blink-timing variability (C) for the X (blue) and AX (orange) tasks across Cue A and Cue B conditions. **(D–F)** Condition means with 95% confidence intervals for the same measures. Circles denote Cue A and squares Cue B. **(G–I)** Parameter estimates from linear mixed-effects models relating blink occurrence (G), mean first-blink latency (H), and blink-timing variability (I) to Task, Cue, and their interaction. Models included overall blink rate, age, sex, and general cognitive ability (*g*) as participant-level covariates. Points represent regression coefficients with 95% confidence intervals; filled symbols indicate significant effects (*p* < 0.05). Experimental predictors are shown above the horizontal divider and participant-level covariates below.

Blink occurrence differed across Task × Cue conditions (Figure 3A, D, G). In the X task, where response selection remained dependent on the target, blink occurrence was similar following A and B cues. In the AX task, where the cue established whether a subsequent target would require a response, blink occurrence was selectively reduced following the A cue, which predicted a task-relevant target with high probability. This differential cue effect was supported by a significant Task × Cue interaction (β = 0.094, 95% CI [0.079, 0.110], *p* <.001; Table 1).

**Table 1.** Linear mixed-effects models predicting blink occurrence, first-blink latency, and blink-timing variability as a function of Task, Cue, their interaction, and participant-level covariates. Regression coefficients (β) are fixed-effect estimates from models including Task, Cue, their interaction, overall blink rate, age, sex, and general cognitive ability (g). Positive coefficients indicate higher values for the second level of each contrast (Task: AX − X; Cue: B − A; Sex: M − F). Values in brackets denote 95% confidence intervals. Significant effects are shown in **bold**.

| *Predictor* | *Blink occurrence* | *First-blink latency* | *Timing variability* |
| --- | --- | --- | --- |
| *Task (AX – X)* | –0.115 [–0.126, –0.104]<br>**< .001** | 0.279 [0.226, 0.332]<br>**< .001** | –0.071 [–0.146, 0.005] .066 |
| *Cue (B – A)* | 0.004 [–0.007, 0.015]<br>.468 | –0.081 [–0.134, –0.028]<br>**.003** | 0.039 [–0.036, 0.115]<br>.308 |
| *Task × Cue* | 0.094 [0.079, 0.110]<br>**< .001** | 0.205 [0.130, 0.280]<br>**< .001** | 0.563 [0.456, 0.670]<br>**< .001** |
| *Blink rate* | 0.225 [0.216, 0.234]<br>**< .001** | –0.653 [–0.724, –0.582]<br>**< .001** | –0.252 [–0.329, –0.176]<br>**< .001** |
| *Age* | 0.003 [–0.006, 0.013]<br>.460 | 0.059 [–0.004, 0.122]<br>.066 | –0.152 [–0.219, –0.084]<br>**< .001** |
| *Sex (M – F)* | –0.006 [–0.025, 0.013]<br>.533 | –0.074 [–0.203, 0.056]<br>.266 | –0.095 [–0.234, 0.044]<br>.180 |
| *Cognitive ability (g)* | 0.005 [–0.004, 0.014]<br>.302 | –0.041 [–0.103, 0.021]<br>.191 | –0.060 [–0.126, 0.007]<br>.077 |

Task context also influenced when participants blinked (Figure 3B, E, H). Mean first-blink latency differed across Task × Cue conditions, with the longest latencies following the B cue in the AX task, where the cue indicated a no-go trial. This pattern was reflected in a significant Task × Cue interaction (β = 0.205, 95% CI [0.130, 0.280], *p* <.001; Table 1).

Blink-timing variability showed a similar context-dependent pattern (Figure 3C, F, I). Variability was highest following the B cue in the AX task, whereas the remaining conditions differed comparatively little. Blink-timing variability likewise showed a significant Task × Cue interaction (β = 0.563, 95% CI [0.456, 0.670], *p* <.001; Table 1).

Across models, participant-level covariates primarily accounted for stable individual differences rather than experimental effects (Table 1). Higher overall blink rates were associated with more frequent, earlier, and less variable blinking, whereas age showed a modest negative association with blink-timing variability. Sex and general cognitive ability were not significantly associated with any blink measure.

Together, these findings indicate that spontaneous blink behavior systematically adapts to task context. Across all three measures, this adaptation depended on the combination of Task and Cue, suggesting that blink behavior is flexibly adjusted to current task demands.

### Individual differences in spontaneous blink behavior are associated with behavioral performance

We next examined whether stable individual differences in blink behavior were associated with behavioral performance. Figure 4 summarizes descriptive associations and mixed-effects models relating blink occurrence, first-blink timing, and blink-timing variability to mean response time.

**Figure 4.**
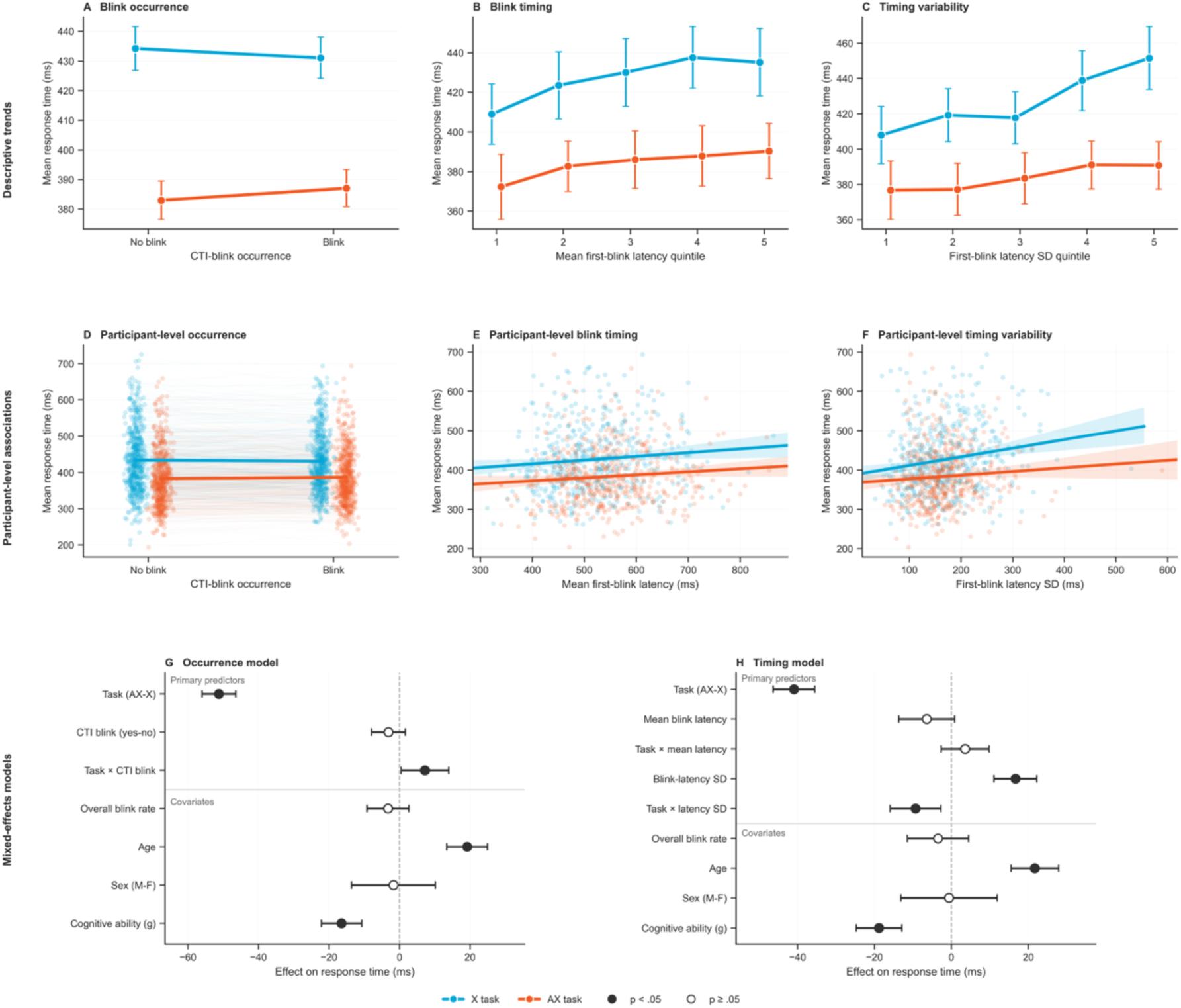
Between-subject associations between spontaneous blink behavior and response time. The figure summarizes descriptive and model-based associations between spontaneous blink behavior and mean response time across participants. **(A–C)** Descriptive associations between blink behavior and mean response time, shown separately for the X (blue) and AX (orange) tasks. Panel A compares participants with and without cue–target interval (CTI) blinks, whereas Panels B and C show mean response time across quintiles of mean first-blink latency and blink-timing variability, respectively. Error bars represent 95% confidence intervals. **(D–F)** Participant-level associations between blink behavior and mean response time. Each point represents one participant in one task condition. Panel D shows within-participant differences between blink-occurrence conditions, whereas Panels E and F show participant-level regression lines with 95% confidence intervals. **(G–H)** Fixed-effect estimates from the covariate-adjusted linear mixed-effects models relating mean response time to blink occurrence (G) and blink timing (H). Models included overall blink rate, age, sex, and general cognitive ability (*g*) as participant-level covariates. Points represent regression coefficients with 95% confidence intervals; filled symbols indicate significant effects (*p* <.05).

Blink occurrence showed only a weak association with behavioral performance (Figure 4A, D, G), with blinking associated with slightly faster responses in the X task but slightly slower responses in the AX task (Task × Blink Occurrence: β = 7.20 ms, 95% CI [0.44, 13.96], *p* =.037; Table 2).

**Table 2.** Linear mixed-effects models predicting mean response time from spontaneous blink behavior. Regression coefficients (β) are fixed-effect estimates from models including task, blink measures, their interactions, overall blink rate, age, sex, and general cognitive ability (g). Positive coefficients indicate higher response times for the second level of each contrast (Task: AX − X; CTI blink: yes − no; Sex: M − F). Values in brackets denote 95% confidence intervals. Significant effects are shown in **bold**.

| *Predictor* | *Blink occurrence $\beta$ (95% CI)* | *p* | *Blink timing $\beta$ (95% CI)* | *p* |
| --- | --- | --- | --- | --- |
| **Primary predictors** |  |  |  |  |
| *Task (AX – X)* | –51.19 [–55.97, –46.41] | **< .001** | –40.94 [–46.33, –35.56] | **< .001** |
| *CTI blink* | –3.15 [–7.93, 1.63] | .197 | — | — |
| *Task × CTI blink* | 7.20 [0.44, 13.96] | **.037** | — | — |
| *Mean blink latency* | — | — | –6.43 [–13.68, 0.81] | .082 |
| *Task × Mean latency* | — | — | 3.56 [–2.67, 9.79] | .262 |
| *Blink-timing variability* | — | — | 16.64 [11.08, 22.20] | **< .001** |
| *Task × Timing variability* | — | — | –9.33 [–15.93, –2.72] | **.006** |
| **Covariates** |  |  |  |  |
| *Overall blink rate* | –3.27 [–9.22, 2.68] | .281 | –3.48 [–11.42, 4.46] | .390 |
| *Age* | 19.16 [13.38, 24.95] | **< .001** | 21.68 [15.49, 27.88] | **< .001** |
| *Sex (M – F)* | –1.72 [–13.61, 10.17] | .777 | –0.62 [–13.16, 11.92] | .923 |
| *Cognitive ability (g)* | –16.44 [–22.13, –10.74] | **< .001** | –18.83 [–24.77, –12.88] | **< .001** |

In contrast, behavioral performance was associated with blink-timing variability but not mean first-blink latency (Figure 4B, C, E, F, H). Participants with greater variability in first-blink latency responded more slowly (β = 16.64 ms, 95% CI [11.08, 22.20], *p* <.001), and this relationship was significantly attenuated in the AX task (Task × Blink-Timing Variability: β = −9.33 ms, 95% CI [−15.93, −2.72], *p* =.006). Mean first-blink latency was not significantly associated with response time (β = −6.43 ms, 95% CI [−13.68, 0.81], *p* =.082), nor did this relationship differ between tasks (Task × Mean Blink Latency: β = 3.56 ms, 95% CI [−2.67, 9.79], *p* =.262).

Across both models, older age was associated with slower response times, whereas higher general cognitive ability (*g*) predicted faster responses (Table 2). Neither overall blink rate nor sex showed significant associations with performance after accounting for the blink measures.

Together, these findings indicate that stable individual differences in the consistency of spontaneous blink timing are behaviorally relevant. Participants with more consistent blink timing responded faster, whereas blink occurrence and mean blink timing showed comparatively little association with performance. We next examined whether this between-participant pattern was complemented by trial-to-trial relationships within individuals.

### Trial-to-trial deviations from individual blink timing

To examine whether the between-participant associations were complemented by within-participant effects, we analyzed trial-to-trial deviations from each participant’s characteristic first-blink latency. Signed deviations quantified whether blinks occurred earlier or later than each participant’s task-specific mean latency, whereas absolute deviations quantified their distance from this characteristic timing irrespective of direction. Nested mixed-effects models tested the contributions of characteristic blink timing, signed and absolute trial-to-trial deviations, and task-dependent effects of these deviations to response time.

Model comparison strongly favored the model including an interaction between Task and absolute blink-timing deviation (Figure 5A). Adding signed deviations produced only modest improvements over the baseline model, whereas absolute deviations substantially improved model fit. Introducing a task-specific effect of absolute deviation further improved model fit, whereas an additional interaction with signed deviation provided no meaningful benefit.

**Figure 5.**
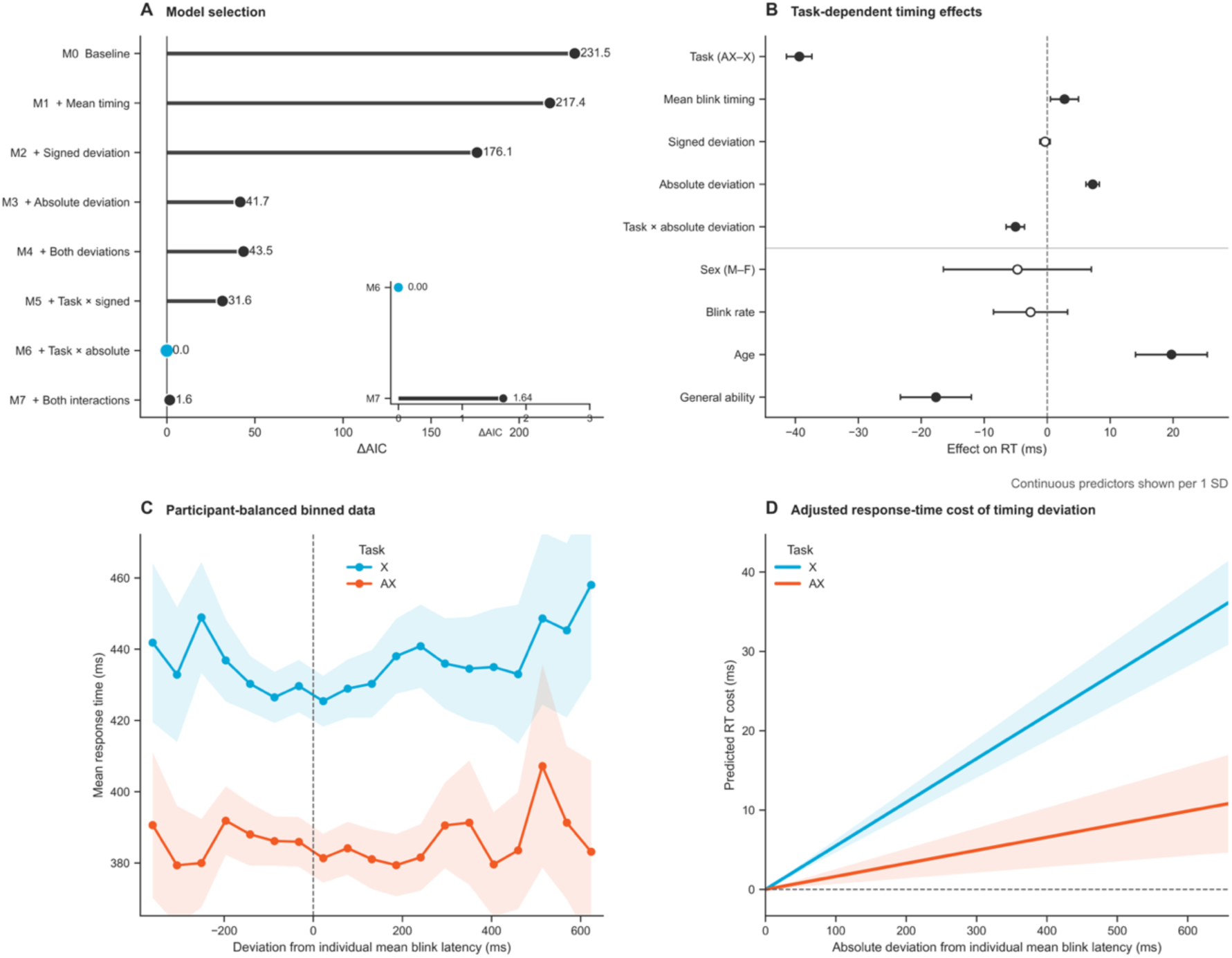
Trial-level analyses of blink timing and response time. (A) Comparison of nested linear mixed-effects models predicting trial-level response time. Models successively included participant-specific mean blink timing, signed and absolute trial-to-trial deviations from each participant’s characteristic blink timing, and their interactions with task. Model fit is expressed as ΔAIC relative to the best-fitting model; the inset highlights the comparison between the two best-supported models. (B) Parameter estimates from the best-fitting adjusted model. Continuous predictors are shown per one standard deviation, whereas task and sex are shown as categorical contrasts. Points represent regression coefficients with 95% confidence intervals; filled symbols indicate significant effects (*p* <.05). (C) Participant-balanced descriptive relationship between signed deviations from each participant’s characteristic blink timing and mean response time. Shaded regions denote 95% confidence intervals. (D) Model-predicted response-time costs as a function of absolute deviation from each participant’s characteristic blink timing, expressed relative to zero deviation while holding all covariates constant. Shaded regions represent 95% confidence intervals.

Parameter estimates from the best-fitting model are shown in Figure 5B. Participants with later characteristic blink timing exhibited slower response times overall. Beyond these between-participant differences, larger trial-to-trial deviations from characteristic blink timing predicted slower responses, with substantially larger performance costs in the X than the AX task.

The descriptive trial-level data mirrored these model-based findings. Response times were fastest for blinks occurring near each participant’s characteristic timing and increased with larger deviations in either temporal direction (Figure 5C). Adjusted predictions from the final model likewise showed an approximately linear increase in response time with absolute deviation, which was markedly steeper in the X than the AX task (Figure 5D).

Together, these findings indicate that the behavioral relevance of blink timing extends beyond stable individual differences. Participants with more consistent blink timing responded faster overall, and responses were fastest on trials in which spontaneous blinks occurred close to each participant’s characteristic timing. Thus, temporal consistency was associated with behavioral performance at both between-and within-participant levels.

## Discussion

The present findings demonstrate that spontaneous blink behavior is systematically organized by task demands and that this temporal organization is meaningfully related to behavior. Across complementary analyses, blink timing adapted to experimentally manipulated task context and predicted both stable individual differences and trial-to-trial fluctuations in response speed. Together, these findings suggest that spontaneous blink timing reflects structured aspects of ongoing cognition rather than merely spontaneous physiology (Murali & Händel, 2021; Pitigoi et al., 2024; Siegle et al., 2008; Wascher et al., 2015).

Task demands influenced whether participants blinked, when blinks occurred, and how consistently they were timed within the cue–target interval. Across cue conditions, blinks occurred later in the AX than the X task. In the X task, cues primarily modulated target expectation, whereas response selection remained dependent on target processing. In the AX task, cues additionally established the response context, likely promoting more extensive cue evaluation and proactive response preparation (Janowich & Cavanagh, 2018; Sharifian et al., 2021).

Cue significance further modulated blink behavior within tasks. In the AX task, A cues were associated with markedly reduced blink occurrence, whereas B cues were followed by later and substantially more variable blinks; cue-related differences in the X task were comparatively small. Thus, the AX task did not impose a general increase in blink suppression but dynamically regulated blinking according to cue significance.

This pattern supports the view that spontaneous blinks are adaptively scheduled according to the changing behavioral relevance of visual information processing (Pitigoi et al., 2024; Wascher et al., 2015). Blink control was strongest while visual information remained behaviorally important and relaxed as processing demands declined, consistent with blinks being scheduled when temporary visual interruption is least costly. This raises the question of which aspects of this adaptive temporal organization are behaviorally meaningful.

The subsequent analyses asked which aspects of blink timing carry behaviorally relevant information. If blink timing indexes the temporal progression of cognitive processing, differences in average blink latency might reflect differences in the duration of underlying cognitive processes, predicting correspondingly earlier or later responses (c.f. Kutas et al., 1977; Polich, 2007; Smulders et al., 1995). The data provided little support for this prediction. Instead, individuals with greater across-trial temporal consistency in blink timing responded faster. Thus, the stability of blink timing across trials, rather than its average temporal position, emerged as the more behaviorally informative characteristic.

The within-participant analyses converged on this conclusion. Incorporating trial-to-trial deviations from characteristic blink timing improved prediction of response time beyond average timing alone. This improvement was driven by deviation magnitude rather than direction: blinks farther from an individual’s characteristic timing were associated with slower responses. Thus, despite their different levels of analysis, the between-and within-participant findings both identify temporal consistency as the behaviorally relevant property of blink timing.

These findings extend evidence that the temporal distribution of blinks contains information beyond average blink latency or blink rate (Ichikawa & Ohira, 2004; Pitigoi et al., 2024). They also parallel broader evidence from cognitive neuroscience that temporal reproducibility of neural dynamics is associated with successful information processing (Churchland et al., 2010; Hanslmayr et al., 2007; Makeig et al., 2002; Mathewson et al., 2009). Within the present paradigm, blink timing may therefore be better understood in terms of the stability with which it remains coupled to the temporal structure of the task than in terms of an optimal average latency.

Temporal consistency may be behaviorally informative for several, non-exclusive reasons. From a mental chronometry perspective, consistent blink timing may reflect stable coupling between spontaneous blinking and the progression of cognitive processing (Murali & Händel, 2021). If blinks align with transitions between processing stages or periods of reduced visual demand, greater regularity of cognitive processing should produce both more consistent blink timing and more stable performance (Ichikawa & Ohira, 2004; Murali & Händel, 2021; Pitigoi et al., 2024). Alternatively, blink timing and performance may be jointly influenced by broader cognitive states. Stable task engagement, attention, or response strategies could support both consistent blinking and performance, whereas fluctuations in vigilance, fatigue, motivation, or effort could disrupt both (Esterman et al., 2014). The present data cannot distinguish these mechanisms. In either case, however, temporal consistency reflects the stability with which blink timing is coupled to the temporal organization of cognition and behavior.

A second, distinct question concerns the source of this temporal organization. Blink behavior was more strongly shaped by cue significance in the AX task, yet its association with behavioral performance was stronger in the X task. Stronger task-driven organization therefore did not translate into a more sensitive readout of individual performance. In the AX task, this organization was particularly evident in the selective suppression of blinking following A cues and its relaxation following B cues. Thus, blink timing appears to reflect not only temporal structure imposed by the task but also temporal organization arising from participant-specific processes.

We propose that the observed chronometric signal reflects the combined influence of these task-imposed and internally generated sources of temporal organization. Task structure constrains when visual interruption is behaviorally costly (Murali & Händel, 2021; Pitigoi et al., 2024; Wascher et al., 2015), whereas participant-specific factors such as mind-wandering, motivation, or fatigue may introduce additional temporal structure or variability (Arnau et al., 2020, 2024; Möckel et al., 2015). These influences may act on blink regulation directly, through the temporal organization of cognitive processing, or both.

From this perspective, stronger cue-dependent regulation in the AX task suggests a greater contribution of task-imposed temporal organization, aligning blink behavior across participants with a common task structure. In the less constrained X task, participant-specific temporal organization may have contributed more strongly to individual variation in blink timing. The stronger blink–response time associations in the X task may therefore reflect greater sensitivity to this participant-specific component rather than greater chronometric sensitivity overall.

The information represented by blink timing may consequently depend on task design. From a mental chronometry perspective, strongly constrained paradigms may be particularly informative about temporal organization imposed by cognitive processing demands, whereas from a neuroergonomic perspective, less constrained paradigms may preserve more information about participant-specific temporal organization related to cognitive state and behavioral performance. These perspectives emphasize complementary aspects of the same chronometric signal rather than competing interpretations. Experimental manipulation of temporal task structure may therefore provide a means of shaping which aspects of temporal organization are expressed in spontaneous blink timing.

The utility of blink timing as a chronometric measure may arise from the unique physiological characteristics of spontaneous blinking. Unlike most physiological processes, spontaneous blinking is neither purely reflexive nor fully voluntary (VanderWerf et al., 2003). Whereas processes such as the heartbeat offer little opportunity for temporal scheduling and deliberate actions are themselves the object of cognitive control, spontaneous blinking occupies an intermediate position: blinks are physiologically inevitable, yet their precise timing remains remarkably flexible (Murali & Händel, 2021; Pitigoi et al., 2024). This unique combination may explain why blink timing is informative. Because blinks cannot simply be omitted but can often be postponed or advanced, their timing reflects the temporal coordination between unavoidable visual interruptions and ongoing cognitive demands, consistent with observations from naturalistic settings (Cornelis et al., 2025; Nishizono et al., 2023).

Rather than treating spontaneous blinks primarily as ocular artifacts requiring removal, the present findings support viewing blink timing as a behavioral chronometric measure whose temporal organization carries information about cognition.

Several limitations should be considered when interpreting the proposed framework. Although the observed associations demonstrate that blink timing reflects structured aspects of ongoing cognition, they do not establish the mechanisms giving rise to this temporal organization. The proposed framework is intended as a conceptual account that generates mechanistic hypotheses for future investigation. In particular, the present data cannot determine the relative contribution of task-imposed temporal structure, the temporal organization of cognitive processing, and processes acting directly on blink regulation to the chronometric information represented by blink timing.

Future research should therefore evaluate blink timing across tasks that systematically vary in temporal structure, cognitive demands, and ecological complexity while developing quantitative models that explain how an unavoidable physiological requirement becomes coordinated with ongoing cognitive processing. Such work will help determine which temporal organization blink timing represents under different task conditions, thereby establishing when and why spontaneous blink timing provides informative chronometric information about cognition.

In conclusion, spontaneous blinking is an unavoidable aspect of human behavior, yet its precise timing remains remarkably flexible. The present findings suggest that this unique combination makes blink timing a valuable behavioral chronometric measure whose interpretation depends on the temporal organization it represents within a given task context. Recognizing this context dependence provides a framework for interpreting spontaneous blink timing across experimental paradigms and for designing tasks that emphasize different aspects of the chronometric information it represents.

## Materials and Methods

### Sample

The present analyses were based on data from the Dortmund Vital Study (ClinicalTrials.gov Identifier: NCT05155397; for a detailed study description, see Gajewski et al., 2023). EEG recordings were initially available from 632 participants. Participants were retained if their EEG recording could be successfully processed, all 480 experimental trials could be reconstructed from valid cue–target event pairs, and complete data on age, sex, and general cognitive ability were available. These criteria were met by 586 participants. An additional 10 participants were excluded during quality control because their overall blink rates fell outside the predefined plausible range. The final sample comprised 576 participants (356 women, 61.8%; 220 men, 38.2%), aged 20–70 years (*M* = 43.81, *SD* = 14.27).

Participants constituted a nonprobability sample recruited through local colleges, companies, and public institutions and through advertisements in newspapers and other public media. All participants provided written informed consent and received €160 for completing the two-day study protocol. The study was approved by the Ethics Committee of the Leibniz Research Centre for Working Environment and Human Factors (Dortmund, Germany) and conducted in accordance with the Declaration of Helsinki.

### Procedure

Participants were seated in a sound-attenuated room in a comfortable armchair. Stimuli were presented on a 32-inch monitor (100 Hz refresh rate), and responses were recorded using force-sensitive keys. The letters A and B served as cues and X and Y as probes. Each trial consisted of a 150-ms cue, an 1850-ms fixation interval, a 150-ms probe, and an 1850-ms response interval. The fixed 2-s cue–target interval was the period of interest for the present blink analyses.

Participants completed an AX Continuous Performance Task (AX-CPT) followed by an X Continuous Performance Task (X-CPT), each comprising 240 trials. Cue–probe combinations AX and BY each occurred on 40% of trials and AY and BX on 10%. In the AX-CPT, participants responded only to X probes preceded by an A cue; in the X-CPT, they responded to every X probe regardless of the preceding cue. Participants were instructed to respond as quickly and accurately as possible.

### Blink Detection

Eye blinks were detected from the continuous EEG using a custom MATLAB algorithm (Alyan et al., 2023a, 2023b), that extends the BLINKER framework (Kleifges et al., 2017) with enhanced signal selection and quality-control procedures. The channel-based detection pathway was used because it provides a standardized signal representation across participants, avoiding variability in the isolation of blink activity by independent component analysis (ICA).

Continuous EEG was band-pass filtered between 1 and 11 Hz. A periorbital composite signal was constructed using Fp1, Fp2, and Fz as the primary electrode set, supplemented by AF7, AF8, AF3, and AF4 when their activity sufficiently corresponded with the primary signal. Candidate blinks were identified using the BLINKER detection procedure and subjected to quality control, including morphological evaluation and rejection of saccade-like events based on bilateral frontal activity. Only events passing all quality-control criteria were retained.

Blinks were assigned to experimental trials when their peak occurred within the cue– target interval (CTI). For each trial, the number of CTI blinks and latency of the first CTI blink relative to cue onset were extracted. Overall blink rate was calculated as the number of accepted blinks per minute across the recording and served as an individual measure of spontaneous blinking.

### Blink Behavior

To characterize blink behavior during the cue–target interval (CTI), two participant-level datasets were constructed from the trial-level master dataset. All trials with valid task and cue information were included irrespective of response accuracy or response time.

The occurrence dataset quantified the probability of blinking during the CTI. For each participant and Task × Cue condition, blink occurrence was calculated as the proportion of trials containing at least one CTI blink. Participants were required to contribute observations to all four conditions.

The timing dataset was restricted to CTI-blink trials with a valid first-blink latency. When multiple blinks occurred, only the first was analyzed, representing the earliest interruption of visual input following cue presentation. For each participant and Task × Cue condition, mean first-blink latency and its within-participant standard deviation were calculated. Each condition was required to contain at least 20 CTI-blink trials, and participants had to meet this criterion in all four conditions. Both timing measures were standardized across participant-by-condition observations before analysis.

Blink occurrence, mean first-blink latency, and first-blink latency variability were analyzed using separate linear mixed-effects models with Task (CPT-X vs. CPT-AX), Cue (A vs. B), and their interaction as fixed effects. Overall blink rate, age, sex, and general cognitive ability (*g*) were included as participant-level covariates, with participant included as a random intercept.

### Blink Timing and Response Time

To examine associations between CTI blinking and subsequent behavioral performance, two participant-level datasets were constructed. Response time was analyzed because accuracy was near ceiling across conditions. Both analyses were restricted to A-cue trials with correct responses and valid, positive response times. A-cue trials provided a comparable response-relevant context across tasks, whereas the behavioral significance of B cues differed between the CPT-X and CPT-AX.

The occurrence dataset examined response time as a function of CTI blink occurrence. Mean response time was calculated for each participant, task, and blink-occurrence condition. Participants were required to contribute data to all four Task × Blink Occurrence conditions.

The timing dataset examined associations between individual differences in blink timing and response time on CTI-blink trials with valid first-blink latencies. For each participant and task, mean response time, mean first-blink latency, and the within-participant standard deviation of first-blink latency were calculated. Participants were required to contribute at least 20 analyzable blink trials in each task and data for both tasks. The two blink-timing measures were standardized across participant-by-task observations.

Mean response time in the occurrence dataset was analyzed using a linear mixed-effects model with Task, Blink Occurrence, and their interaction as fixed effects. Mean response time in the timing dataset was analyzed using a separate model with Task, standardized mean first-blink latency, standardized first-blink latency variability, and their interactions with Task. Both models included overall blink rate, age, sex, and general cognitive ability (*g*) as participant-level covariates and participant as a random intercept. Continuous covariates were standardized across participants.

### Within-Subject Deviations in Blink Timing

To complement the between-participant analyses, we examined whether trial-to-trial deviations from an individual’s characteristic blink timing predicted response time. If blink timing is adapted to task demands, deviations from characteristic timing should be associated with longer response times.

The analysis included A-cue trials with correct responses, valid response times, and a valid first CTI blink. Participants were required to contribute at least 10 analyzable blink trials per task. Characteristic blink timing was defined as each participant’s mean first-blink latency within each task. Signed deviation was calculated as the difference between trial-specific and characteristic blink latency, with negative and positive values indicating earlier-and later-than-usual blinks, respectively. Absolute deviation quantified the magnitude of this difference irrespective of direction.

Nested linear mixed-effects models were used to determine whether increasingly specific representations of blink timing improved prediction of response time. Starting from a baseline model including Task, overall blink rate, age, sex, and general cognitive ability (*g*), successive models added characteristic blink timing, trial-specific deviations, and task-dependent effects of these deviations. Continuous participant-level covariates were standardized across participants, and participant was included as a random intercept. All models were fitted to the same complete-case sample and compared using Akaike’s Information Criterion and likelihood-ratio tests. The best-fitting model was used to estimate blink-timing effects on response time.

## Code and data availability

The code used to analyze the data and creating the figures can be found at https://github.com/stefanarnau/cpt_blink_latencies. The blink data is available at https://osf.io/7zp3r/overview (will be made public with publication).

## Author Contributions

**Stefan Arnau:** Conceptualization, Formal Analysis, Methodology, Visualization, Writing – Original Draft. **Anna Plachti:** Conceptualization, Writing – Review & Editing. **Emad Alyan:** Methodology, Writing – Review & Editing. **Edmund Wascher:** Conceptualization, Writing – Review & Editing. **Daniel Schneider:** Conceptualization, Methodology, Writing – Review & Editing.

